# Scaling SMILES-based chemical language models for therapeutic peptide engineering

**DOI:** 10.64898/2026.01.06.697994

**Authors:** Aaron L. Feller, Maxim Secor, Sebastian Swanson, Claus O. Wilke, Kristine Deibler

## Abstract

Therapeutic peptides occupy a unique middle ground in drug discovery, offering the high specificity of protein interactions with the chemical diversity of small molecules, yet they currently fall in a computational blind spot. Existing foundation models cannot handle them effectively: protein models are restricted to natural amino acids, while chemical models struggle to process large, polymer-like sequences. This disconnect has forced the field to rely on static chemical descriptors that fail to capture subtle chemical details or on complex multi-embedding pipelines that are custom tailored to specific datasets. To bridge this gap, we present PeptideCLM-2, a suite of chemical language models trained on over 100 million molecules to natively represent complex peptide chemistry. This modeling approach expands the available toolkit of machine learning models for therapeutic peptides. Benchmarking results show strong performance versus prior methods for predicting development endpoints including membrane diffusion, biological function, and half life.

## Introduction

Peptides are a rapidly evolving modality in the therapeutic space. ^1–6^ They occupy a unique chemical niche situated between small molecules and proteins. Like small molecules, they possess immense chemical diversity.^7–11^ Yet, they retain the modularity of biological polymers, allowing for accessible and efficient synthesis.^12–14^ While modern synthetic methods enable the creation of a vast diversity of modified peptides, the computational tools required to rationally select among these options have not kept pace. Attempts to adapt existing frameworks have faced limitations. Protein language models (pLMs) are restricted to fixed amino-acid alphabets, rendering them unable to encode noncanonical or chemically modified residues.^15^ Conversely, chemical language models (CLMs) are typically trained on small molecules, lacking the contextual range to interpret peptide-specific motifs.^16,17^

Large-scale protein corpora such as UniProt^18^ have fueled the rise of self-supervised pLMs. Architectures like the ESM family^19–21^ and ProtTrans^22^ have proven that transformers can learn structural and functional constraints directly from sequence data. These models capture deep dependencies predictive of structure,^23–25^ variant effects,^26,27^ and molecular function.^28,29^ In parallel, CLMs utilizing symbolic representations like SMILES or SELFIES^30,31^ have enabled encoding of chemical syntax via masked or autoregressive objectives,^32–34^ enabling prediction of various chemical properties.^35,36^ Extensions such as ChemBERTa-2^37^ and ProLLaMA^38^ have further suggested that augmenting self-supervision with physicochemical regression introduces a valuable inductive bias, improving generalization. Predict ive frameworks for noncanonical peptides are evolving rapidly. Recent methodologies span diverse strategies including structural prediction,^39,40^ message-passing networks such as ChemProp,^41,42^ graph transformers,^43^ and multi-feature representations.^44^ Despite these advances, deep learning has shown minimal improvement over molecular fingerprints for peptide property prediction.^45,46^

In this work, we introduce PeptideCLM-2, a suite of nine pretrained SMILES transformer encoders^47^ designed to excel at modeling modified peptides. Building upon our previous architecture, PeptideCLM,^48^ we train models across a range of 32 to 337 million parameters with three distinct objectives: masked language modeling (MLM), multi-task regression^49^ (MTR) to RDKit-derived descriptors,^50^ and a dual objective combining both. This systematic design provides a framework to rigorously decouple the effects of model scale and pretraining methods on representation learning. As therapeutic peptides often adopt transient rather than static conformations, we employed a string-based architecture to capture topological connectivity without the bias of a single, rigid 3D structure.

What follows is a rigorous evaluation of the models released here, showing that these models perform competitively with molecular fingerprints, structure-based modeling, alternative BERT encoders, and specialized architectures. Our evaluation reveals that molecular descriptors are competitive for prediction on simple tasks, while highly similar molecules provide a more difficult problem that deep learning excels at. These results establish Pep-tideCLM-2 as both an open, scalable resource for peptide engineering and a framework for understanding the interaction between model capacity and pretraining for the field of computational peptide chemistry.

## Results

### Transformer architecture and multi-objective pretraining for diverse peptide chemistries

The PeptideCLM-2 suite represents a unified framework for embedding therapeutic peptides. Unlike standard protein language models, which are restricted to a fixed alphabet of 20 canonical amino acids, PeptideCLM-2 is able to encode all of chemical space. By combining scalable transformer architectures with interpretable tokenization strategies, the framework provides a bridge between symbolic chemical representation and chemical function.

To rigorously evaluate chemical deep learning for peptides, we designed a controlled grid of nine models with varied model capacity across three orders of magnitude—32M, 114M, and 337M parameters—and trained each scale using three distinct pretraining objectives: masked-language modeling (MLM), multi-task regression (MTR) to physicochemical descriptors, and a combined dual objective. This systematic design provides insight into the effects of parameter scale and pretraining methods.

PeptideCLM2 processes inputs as raw SMILES strings, allowing for input to contain canonical residues, noncanonical modifications, cyclic scaffolds, and complex conjugations like lipidation or PEGylation (Fig. 1A). The backbone of the framework is a BERT-style transformer encoder (Fig. 1B). To improve training stability and effective handling of long-range chemical dependencies over prior work, ^48^ we incorporated modern architectural features including rotary positional embeddings,^51^ SwiGLU activation functions,^52^ and pre-layer normalization.^53^ Model depth and hidden dimension scale with parameter count, maintaining a consistent per-head dimensionality of 64.

**Figure 1:**
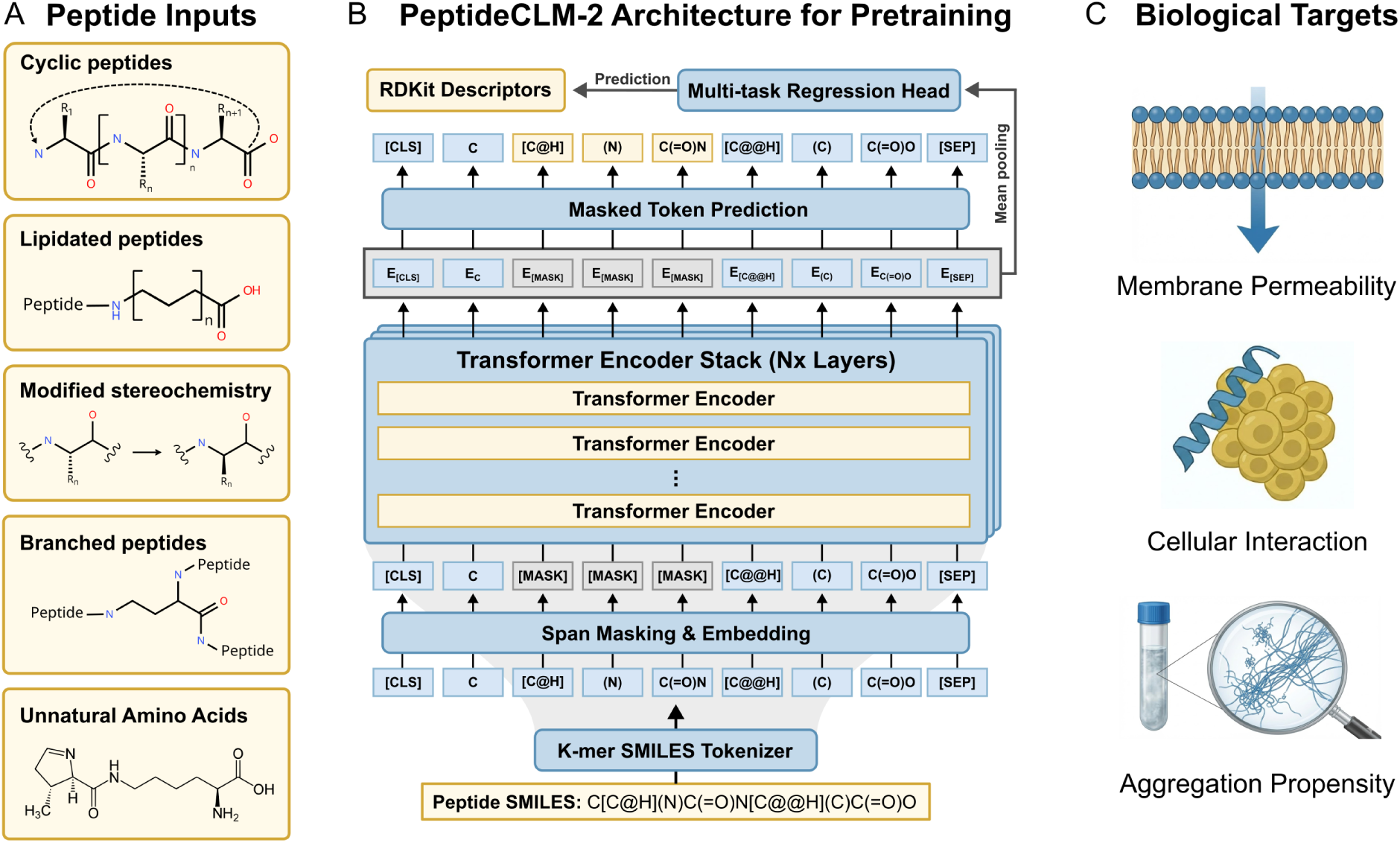
PeptideCLM-2 workflow: enabling encoding of chemical diversity for predictive modeling. (A) The model can input SMILES strings from peptides with various modifications, allowing it to encode cyclic peptides, noncanonical amino acids, and synthetic modifications. (B) Models were trained with both span masking (predicting grey [MASK] tokens) and regression to physicochemical descriptors (e.g., LogP, TPSA) from a pooled sequence embedding. (C) Examples of prediction targets for downstream tasks, including membrane permeability, cellular interactions, and aggregation propensity.

We investigated three distinct learning paradigms to train the architecture. The masked language modeling (MLM) objective utilized span masking^54^ to force the model to reconstruct missing chemical fragments, effectively learning the syntax of molecular graphs. The multi-task regression (MTR) objective explicitly grounded the embeddings with regression to 99 RDKit-derived physicochemical descriptors (Table S1) from a mean-pooled embedding; the transformer itself still receives only SMILES strings at inference time. Finally, a hybrid objective optimized both losses simultaneously, testing whether syntax and semantics could be learned in tandem. For downstream evaluation, the pretrained encoder is coupled with a lightweight, 2-hidden layer feed-forward regression head employing GeLU activation, allowing the model to adapt its learned representations to specific biological tasks such as membrane permeability, cellular interactions, or aggregation (Fig. 1C).

### Composite pretraining corpus and sequence compression via k-mer tokenization

To construct a representation space that spans the full continuum from small molecules to biological polymers, we curated a composite pretraining corpus combining three distinct datasets. We aggregated lipids from the LIPID MAPS Structure Database (*n* = 49, 145),^55^ small drug-like molecules from PubChem (*n* = 108, 583, 157),^56^ and diverse peptide sequences from ESMAtlas (*n* = 9, 634, 945). ^20^ These datasets provide distinct regions of chemical space (Fig. 2A), ensuring the model encounters a heterogeneous distribution of molecular syntax.^57^ Due to the flexibility of SMILES encoding, we are able to train on small molecules to learn chemical features, which can directly translate to therapeutic peptides that contain chemistries not seen in the natural amino acid space.

**Figure 2:**
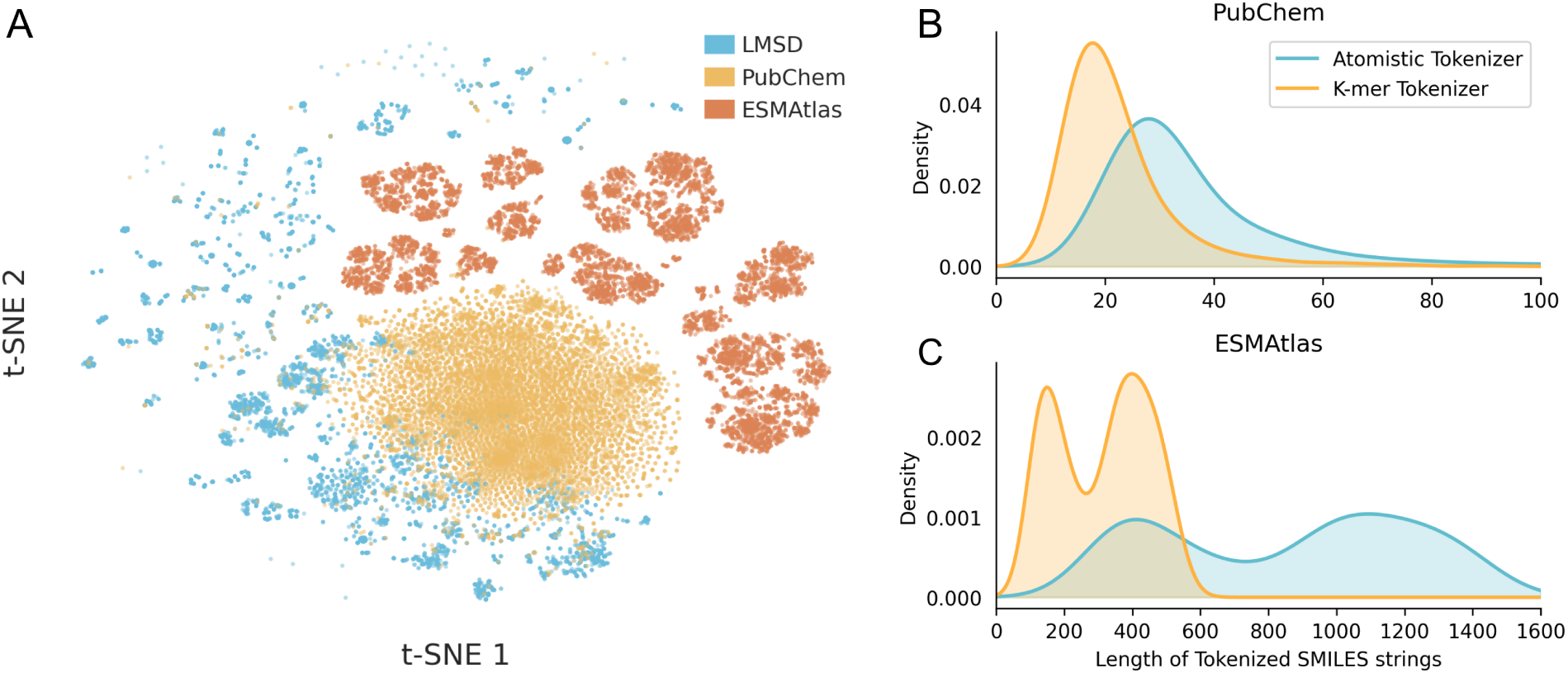
Data Diversity and Representational Efficiency. (A) A 2D t-SNE projection of Morgan Fingerprints from the pretraining corpus, demonstrating that the composite dataset (LMSD, PubChem, and ESMAtlas) bridges distinct chemical subspaces. Comparative length distributions of tokenized molecules shows k-mer tokenization strategy yields a substantial compression factor relative to atom-level encoding, reducing sequence length by (B) 38% for small molecules and (C) 64% for peptides.

A central bottleneck in applying chemical language models to biopolymers is the computational cost of self-attention, which scales quadratically with sequence length (O(*n*^2^)). Because direct character encoding of peptides generates exceptionally long SMILES strings, standard models become prohibitively expensive to train. To resolve this, we developed a specialized k-mer tokenizer (Tables S2 and S3) that compresses these symbolic representations by mapping recurring sub-structural motifs to single tokens. This strategy reduces the effective sequence length, and thus the quadratic compute burden, while maintaining full compatibility with standard SMILES syntax.

Quantitatively, this strategy reduces sequence lengths by 38% for small molecules (Fig. 2B) and 64% for natural peptides (Fig. 2C) relative to standard atom-level frameworks like DeepChem.^58^ In order to conduct a fair evaluation, we held the released pretraining configuration constant across model-scale experiments (Table S4). We similarly standardized downstream workflows and optimization settings across finetuning experiments (Tables S5 and S6).

We found that this compression comes at no cost to accuracy; benchmarking for the small model size resulted in equivalent performance on membrane permeability prediction for models trained with either tokenizer. Thus, the k-mer approach successfully resolves the trade-off between computational tractability and semantic fidelity, allowing the model to attend to long-range dependencies in complex backbones without the prohibitive cost of character-level encoding.

### Parameter scaling drives increased performance on prediction of membrane permeability

We evaluated PeptideCLM-2 using the CycPeptMPDB dataset,^59^ a standard benchmark for cyclic peptide permeability with robust evaluation splits.^48^ To ensure a rigorous comparison, we focused our analysis on nine distinct model configurations selected via ablation studies, covering three distinct parameter sizes (Table S7).

We first investigated whether the models had learned a chemically meaningful latent space. By projecting the embeddings of the 337M MLM model into two dimensions, we observed how the model organizes the chemical manifold and how that relates to physicochemical properties. When visualizing embeddings from the large PeptideCLM-2 model trained with MLM-only, we can see some separation across molecules by molecular weight (Fig. 3A, B), aromaticity (Fig. 3B), and other physicochemical descriptors including charge, logP, and available hydrogen bonding sites (Fig. S1). The projection of high-dimensional embeddings shows some coarse clustering but no overall organization of measured PAMPA permeability (Fig. 3C).

**Figure 3:**
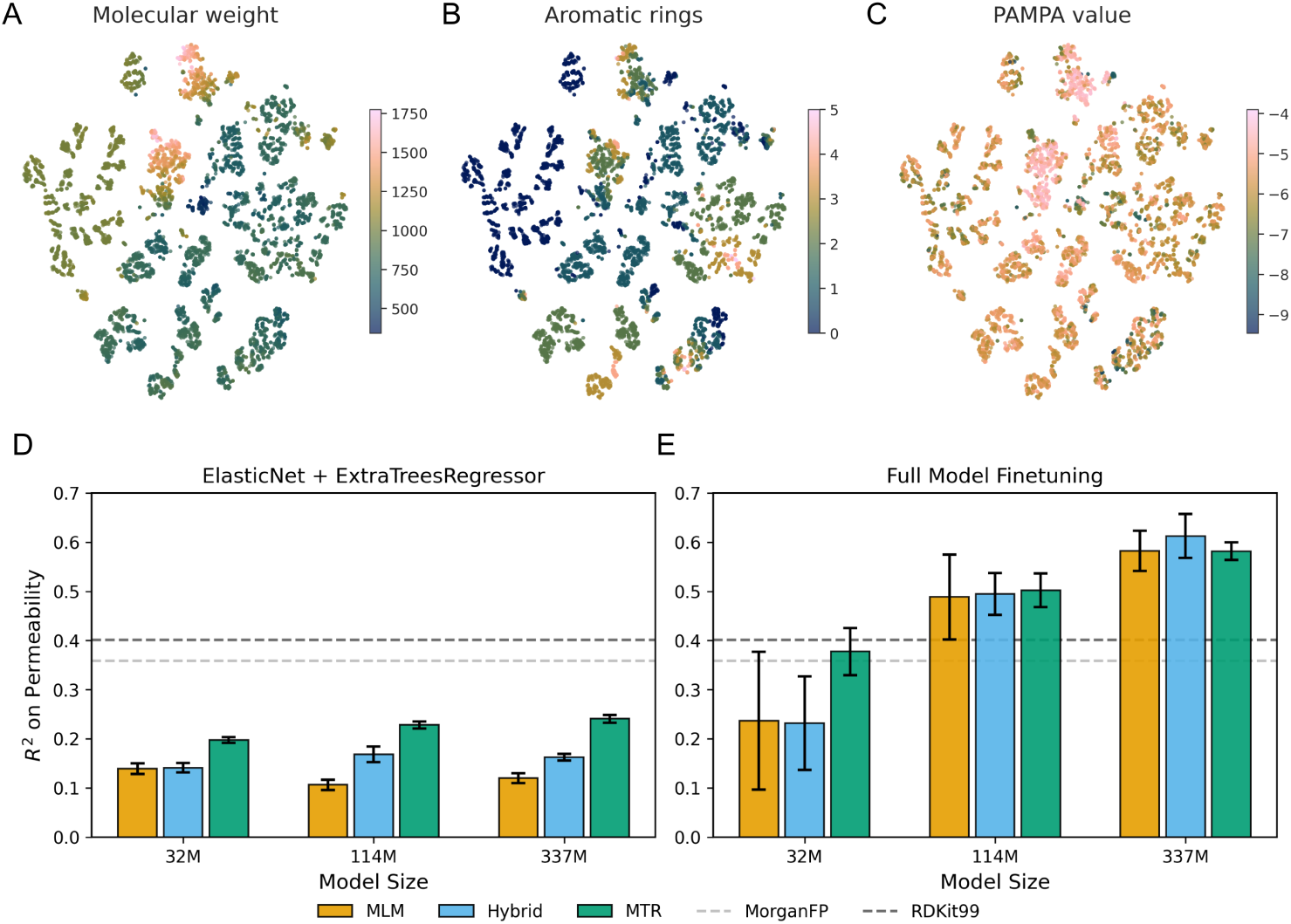
Model embeddings are loosely correlated with chemical properties and model scale improves predictive capability. (A-C) t-SNE projections of embeddings from PeptideCLM-2 337M parameter MLM model. The points are colored by (A) molecular weight, (B) aromaticity, and (C) measured permeability. (D) *R*^2^ scores for nine PeptideCLM-2 model variants using final-layer mean-pooled embeddings trained with a 50/50 ensemble of ElasticNet and ExtraTreesRegressor. Dashed lines represent Morgan fingerprints (MorganFP) and 99 RDKit descriptors (RDKit99). (E) Full model finetuning with all weights active with increasing *R*^2^ as model size scales.

To disentangle the quality of these raw representations from the model’s capacity to adapt, we compared downstream performance using frozen model embeddings against full-model finetuning. When utilizing a 50/50 ensemble of ElasticNet^60^ and ExtraTreesRegressor^61^ trained on frozen features, predictive performance was generally limited (*R*^2^ *<* 0.30) regardless of model scale, though MTR pretraining consistently yielded the highest baseline performance (Fig. 3D). In these comparisons, MorganFP serves as a topology-only baseline, whereas RDKit99 serves as a descriptor-only baseline aligned with the MTR supervision target. Full-model finetuning revealed a dependency on the pretraining strategy at lower parameter scales. Specifically, the 32M parameter model pretrained with explicit physicochemical regression (MTR) achieved a higher mean test score and a smaller run-to-run standard deviation than its MLM and Hybrid counterparts *R*_MTR_^2^ = 0.38 ± 0.05 vs. *R*_Hybrid_^2^ = 0.23 ± 0.10 vs. *R*_MTR_^2^ = 0.24 ± 0.14) This indicates that lower-capacity models may benefit from pretraining that is grounded in physical properties. As model capacity increased to 114M and 337M parameters, this dependency on the pretraining method became negligible, where the SD for all three pretraining modalities overlapped.

To explore how information is propagated through the model, we probed each layer of the network with a linear model. Training on pooled embeddings from each layer revealed a difference in stability between model sizes. In smaller models, the final hidden layer—typically used for downstream tasks—often underperformed compared to intermediate layers (Fig. S2), suggesting a forgetting of physicochemical details at the output bottleneck. In contrast, larger models exhibited high stability, maintaining rich representations from the middle layers through to the final output.

### Predictive generalization to complex biological phenotypes and non-canonical chemistry

We evaluated the large-sized models on three diverse classification benchmarks—tumor homing, cell penetration, and antimicrobial activity (Tables S8–S10). The datasets used for these tasks all include noncanonical chemistry. Embedding projections from the PeptideCLM-2 MLM 337M parameter model show distinct spatial separation between positive and negative classes across all three benchmarks (Fig. 4A, C, E). This latent structure provides a robust foundation for downstream prediction, where PeptideCLM-2 models consistently met or outperformed specialized architectures explicitly designed to encode modified peptides.

**Figure 4:**
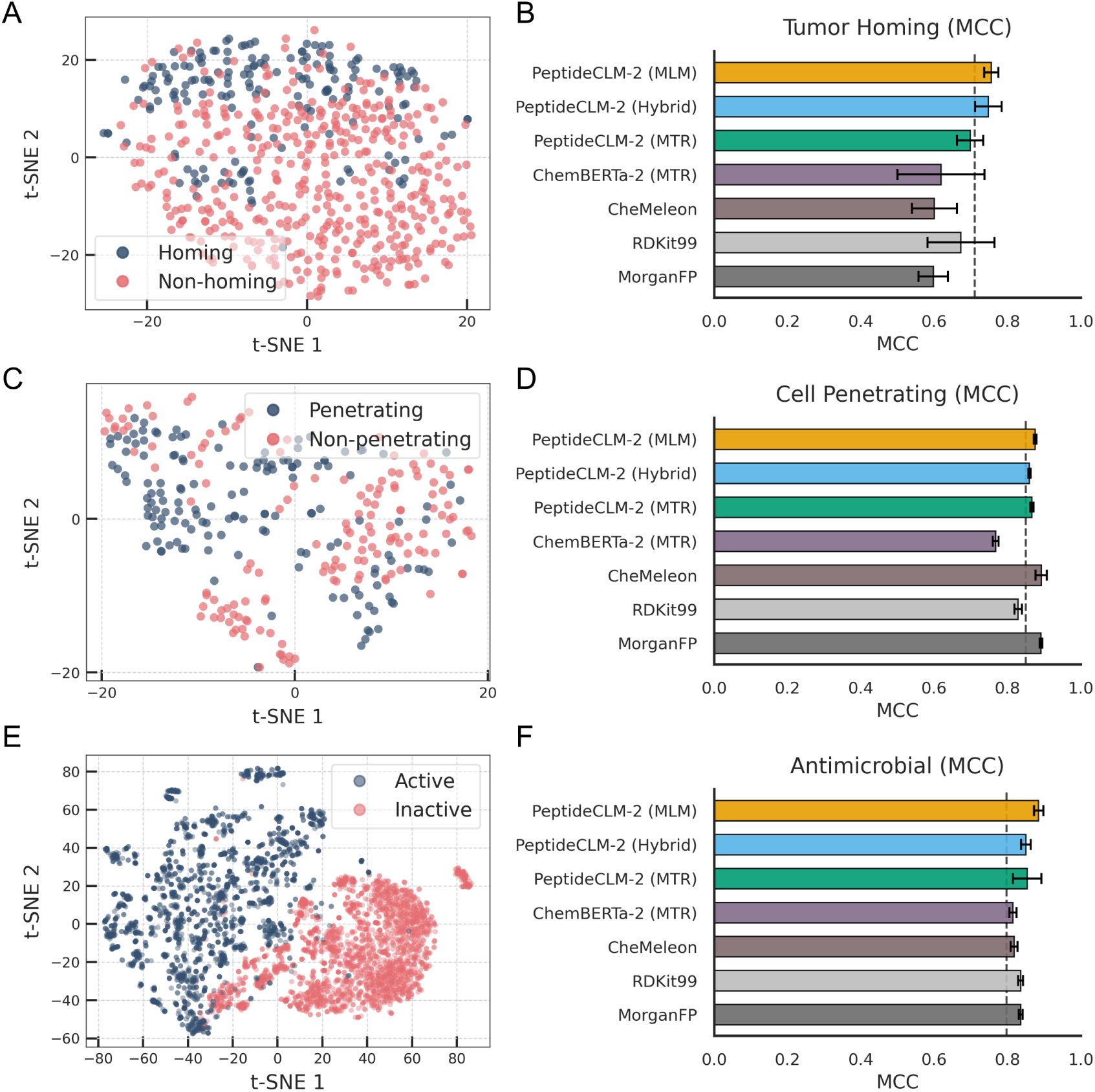
Model evaluation using three benchmark datasets of biological interaction. Left panels: t-SNE projections (A, C, E) of embeddings from the 337M Hybrid model. Even before finetuning, the model’s latent space exhibits strong linear separability between active (blue) and inactive (red) peptides across all three tasks. **Right panels:** Performance comparison (MCC) between the three PeptideCLM-2 Large models, ChemBERTa-2 77M MTR, CheMeleon, and XGBoost models trained on RDKit99 and MorganFP (B, D, F). Grey dashed lines indicate the performance of prior state-of-the-art methods (THPep, ^62^ CellPPD,^63^ AmpHGT^64^).

#### Tumor Homing

Tumor-homing peptides rely on subtle recognition motifs (e.g., RGD, NGR) to target tumor vasculature.^65^ The baseline method published with this dataset, TH-Pep, achieved an MCC of ≈ 0.71 by relying on hand-engineered features, such as Pseudo Amino Acid Composition (PseAAC).^62^ PeptideCLM-2 surpassed this baseline (MCC = 0.756) using only raw SMILES strings (Fig. 4B), demonstrating that the model effectively learns motif-driven features without explicit feature engineering.

#### Cell Penetration

On the CellPPD benchmark^63^—a dataset explicitly containing chemically modified peptides—prior modeling efforts relied on computing extensive 2D/3D chemical descriptors via PaDEL^66^ to capture side-chain modifications. In contrast, PeptideCLM-2 encodes the entire modified macrocycle in a single pass directly from chemical syntax, achieving an MCC of 0.875 and outperforming the original CellPPD baseline (≈ 0.85; Fig. 4D).

#### Antimicrobial Activity

A challenging benchmark designed by He et al.^64^ evaluates out-of-distribution generalization by training models strictly on natural peptide sequences and testing performance on peptides containing noncanonical amino acids. The specialized baseline, AmpHGT, employs a complex Heterogeneous Graph Transformer to explicitly map atoms and fragments, achieving an MCC of 0.797. PeptideCLM-2 surpassed this graph-based architecture with an MCC of 0.884 (Fig. 4F).

Together, these benchmarks highlight a critical architectural advantage of PeptideCLM-2: its capacity to natively reconcile the distinct vocabularies of biological sequences and raw chemistry. While prior methods required heavily engineered features (THPep), specialized side-chain descriptor parsing (CellPPD), or explicit atom-fragment graph mappings (AmpHGT), a unified chemical language approach effectively internalizes these structural constraints. The model’s superior performance on the antimicrobial split confirms that its learned representations do not merely rely on sequence-based memorization, but instead capture transferable physicochemical features that generalize robustly to entirely unseen noncanonical spaces.

### Prediction of higher order peptide properties evaluated via half-life and aggregation propensity

Peptide stability dictates therapeutic viability through two primary mechanisms: resistance to enzymatic degradation in circulation and the absence of aggregation during storage. We evaluated PeptideCLM-2 on both axes to determine if the learned chemical syntax successfully captures these complex biophysical properties.

To assess enzymatic stability, we utilized the recently curated PepMSND dataset,^67^ a benchmark of peptide half-life spanning 635 diverse samples evaluated across varying blood environments. The originally published modeling approach for this dataset relied on a complex multimodal ensemble combining 0D physicochemical descriptors, 1D sequences, 2D molecular graphs, and 3D structural conformations. This baseline incorporates a Kolmogorov-Arnold Network^68^ (KAN) to process tabular molecular descriptors, which ablation studies^67^ identified as the largest contributor to overall predictive capability.

Visualizing the pretrained PeptideCLM-2 embeddings for the PepMSND dataset (Fig. 5A) reveals several local clusters dominated by a single class (stable vs. unstable), though a substantial portion of the latent landscape is a blend of the two classes. Despite this visual complexity, PeptideCLM-2 outperforms the baseline multimodal approach^67^ when the latter lacks its descriptor KAN component (Fig. 5B). This demonstrates that large-scale representation learning can effectively substitute for intricate, multi-level feature engineering in blood stability workflows. Furthermore, combining PeptideCLM-2 embeddings with the same KAN architecture and feature-fusion regression strategy implemented in PepMSND^67^ yields a performance lift that surpasses the original state-of-the-art PepMSND+KAN baseline (Table S11).

**Figure 5:**
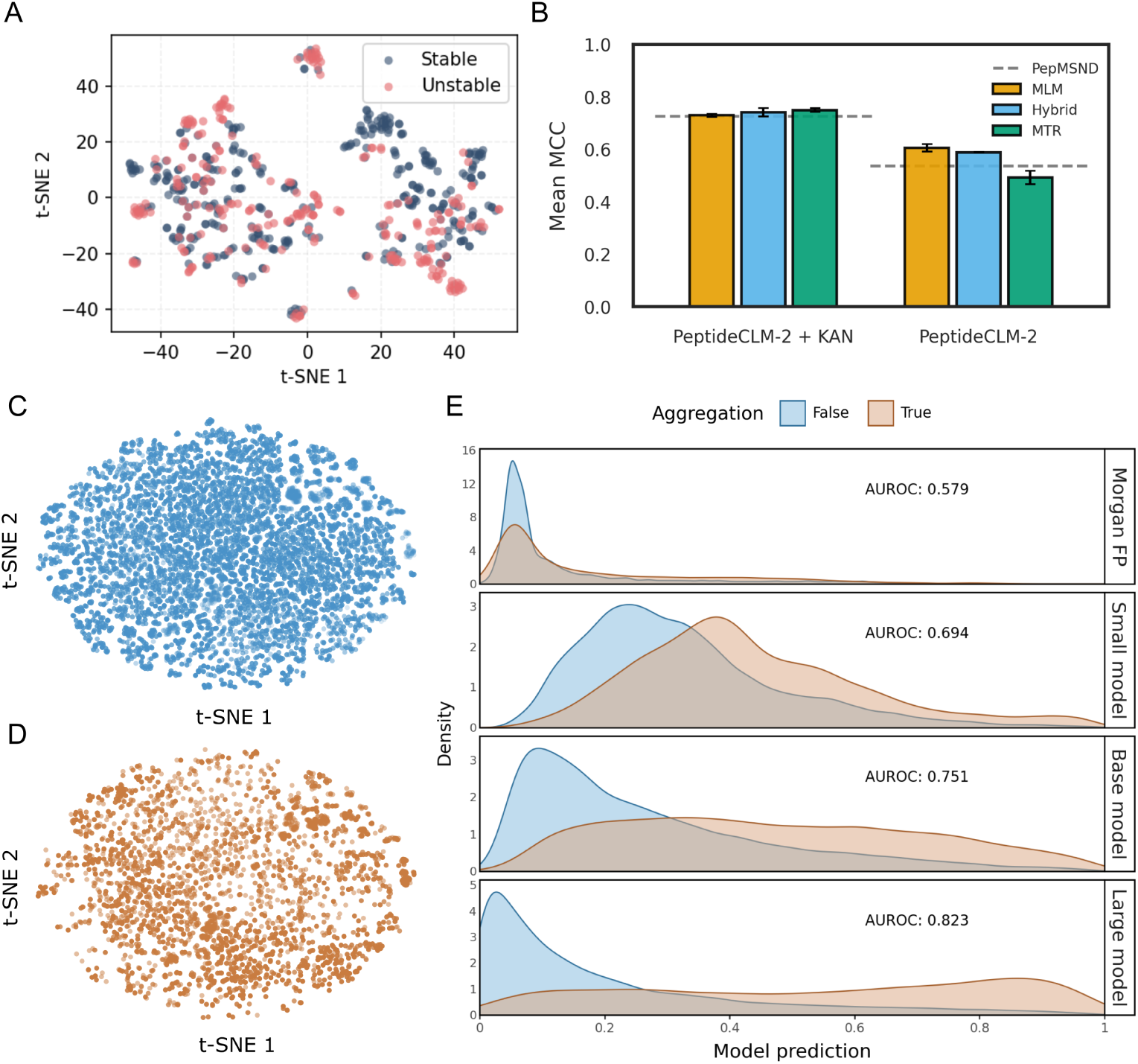
Scaling model size in predicting peptide stability. (A) t-SNE projection of PeptideCLM-2 MLM model embeddings for all peptides present in PepMSND, colored by either a passing or failing stability checkpoint. (B) Bar chart of PeptideCLM-2 predictions on binary stability, reported as the mean ± standard deviation across three independent random seeds evaluated on the PepMSND test splits, colored by model type. KAN represents a Kolmogorov-Arnold Network trained on molecular descriptors, as used in the original PepMSND pipeline. The grey dashed line indicates the baseline model score from the original publication. t-SNE visualizations of the peptide fibrillation data show overlap between non-aggregating (C) and aggregating (D) peptides, illustrating the difficulty of linearly separating these classes based on structure alone. (E) Density plots of model predictions across the two fibrillation populations. Traditional molecular fingerprints (Morgan FP) achieve an AUROC of 0.579, whereas PeptideCLM-2 performance scales systematically with model capacity, reaching an AUROC of 0.823 at the 337M parameter scale. Density curves are scaled independently for visibility.

Beyond enzymatic degradation, physical aggregation represents a critical failure mode driven by condition-dependent interactions with lipids, excipients, and secondary structure shifts. To evaluate prediction with of this phenomenon with PeptideCLM-2, we utilized a large proprietary dataset consisting of Thioflavin T (ThT) fluorescence assays. This benchmark encompasses engineered macrocycles and endogenous peptides containing diverse protraction moieties measured across varying pH conditions, with a roughly 2-to-1 data imbalance favoring non-aggregating peptides. To allow the network to learn these environment-dependent stability profiles, the experimental pH value was appended directly to the embedding vector prior to the regression head.

The inherent difficulty of predicting fibrillation is immediately apparent in the latent space. The embedding projections exhibit extensive coordinate overlap between the aggregating and non-aggregating peptide populations (Fig. 5C, D). This spatial overlap is primarily driven by the fact that identical molecular structures evaluated under differing assay pH conditions naturally map to identical initial structural coordinates. A baseline Random Forest model trained strictly on Morgan fingerprints and pH values achieved an area under the receiver operating characteristic curve (AUROC) of just 0.579, confirming that standard topochemical features and pH descriptors do not provide sufficient class separation.

In contrast, PeptideCLM-2 demonstrates a robust recovery of predictive power that scales positively with expanded model capacity (Fig. 5E). While the 32M parameter model establishes a baseline discrimination layer (AUROC = 0.694), scaling to the 114M and 337M parameter architectures systematically improves performance to 0.751 and 0.823, respectively. This distinct scaling trajectory confirms that large chemical language models can capture the subtle, non-linear biophysical drivers of aggregation that remain invisible to static chemical fingerprints.

### Summary of predictive performance

A comprehensive evaluation across six distinct datasets demonstrates competitive performance of PeptideCLM-2 as compared to prior methods,^48,62–64,67^ molecular descriptors,^50,57^ CheMeleon,^42^ and ChemBERTa-2.^37^ All datasets evaluate feature peptides incorporating noncanonical amino acids and complex chemical modifications that traditional protein language models cannot process. Notably, XGBoost models trained on RDKit descriptors or Morgan Fingerprints performed remarkably well. For prior methods, PeptideCLM-2 out-performs the models published alongside the datasets and training splits used in this study (Table 1). Despite the competitive landscape for model architectures and prediction approaches, having a transformer encoder with competitive performance opens the potential for various applications including cross-attention, steering, and a general advance in one modeling approach.

**Table 1:** Summary of performance across five therapeutic peptide benchmark datasets. Values represent mean score ± standard deviation across independent seeds. Bold values indicate top performance among evaluated models, while underlined values denote second place within each task column.

| Model | Perm.<br>(AUROC) | THP<br>(MCC) | CPP<br>(MCC) | AMP<br>(MCC) | Stab.<br>(MCC) |
| --- | --- | --- | --- | --- | --- |
| *Prior Published Baselines* |  |  |  |  |  |
| Alt. Baseline | PeptideCLM<sup>48</sup> | THPep<sup>62</sup> | CellPPD-Mod<sup>63</sup> | AmpHGT<sup>64</sup> | PepMSND<sup>67</sup> |
| Alt. Score | 0.781 | 0.710 | 0.850 | 0.797 | 0.726 |
| *Baseline Implementations* |  |  |  |  |  |
| MorganFP | 0.792±0.001 | 0.597±.040 | <u>0.891</u>±.003 | 0.836±.005 | — |
| RDKit99 | 0.800±0.003 | 0.673±.092 | 0.829±.010 | 0.835±.007 | — |
| *Competitive Models* |  |  |  |  |  |
| CheMeleon | — | 0.600±.062 | **0.892**±.016 | 0.818±.009 | — |
| ChemBERTa-2 (MTR) | — | 0.618±.118 | 0.768±.008 | 0.814±.010 | — |
| *Our large models* |  |  |  |  |  |
| PeptideCLM-2 (MTR) | 0.825±.009 | 0.698±.036 | 0.867±.004 | <u>0.853</u>±.039 | **0.749**±.007 |
| PeptideCLM-2 (Hybrid) | **0.836**±.011 | <u>0.747</u>±.036 | 0.860±.003 | 0.850±.014 | <u>0.741</u>±.016 |
| PeptideCLM-2 (MLM) | <u>0.829</u>±.020 | **0.756**±.019 | 0.875±.004 | **0.884**±.013 | 0.729±.006 |

**Task Details:** **Perm.:** Membrane Permeability (PAMPA dataset); **THP:** Tumor Homing Peptide; **CPP:** Cell Penetrating Peptide; **AMP:** Antimicrobial Activity; **Stab.:** Blood Stability (PepMSND dataset benchmark).

## Discussion

In-silico analysis of therapeutic peptides is hindered by a representational dilemma. While machine learning has revolutionized protein engineering, peptides remain stranded between two paradigms: protein language models cannot handle noncanonical chemistry, while chemical language models lack the contextual depth to model long biopolymers. Our initial foray into this space, PeptideCLM,^48^ demonstrated the feasibility of SMILES-based modeling for cyclic peptides. In this work, we expand this concept into a comprehensive framework, Pep-tideCLM-2, incorporating advances in data scale, architectural depth, and rigorous bench-marking that demonstrate the utility of transformer-based encoders for modified peptides.

Our analysis presents lessons for effective peptide modeling. First, we introduce a novel k-mer tokenizer to process these complex molecules efficiently. This innovation allows for shorter encoding length, thus allowing further scaling to greater parameter models, pushing toward improved results. Second, we observe a parameter scaling effect where larger models have greater performance when finetuned on labeled data. Finally, our results demonstrate that supervised pretraining predicting calculable molecular descriptors improves predictive capability on small scale models, a property that diminishes at scale where self-supervised modeling is competitive.

### Large-scale transformers spontaneously derive physical rules from chemical syntax alone

Our results demonstrate that the optimal pretraining strategy is strictly a function of model capacity. At the 32M parameter scale, it seems that the model lacks the representational capacity to infer thermodynamic laws from raw syntax alone. In this regime, explicit physico-chemical supervision provides a decisive advantage, acting as a scaffold that guides the model toward a structured embedding space where permeability can be accurately predicted.

However, this advantage vanishes at scale. The convergence of self-supervised and descriptor-guided models at 337M parameters confirms that sufficiently large transformers outperform chemical features when trained only on token cooccurrence. This finding offers a mechanistic explanation for prior observations in the field, such as the reported superiority of descriptor-guided training in ChemBERTa-2.^37^ We posit that ChemBERTa-2, being a relatively lightweight architecture, operated on the lower end of this scaling curve, where inductive bias is still beneficial, while MoLFormer,^34^ trained purely on SMILES, operated on the upper end of this curve. PeptideCLM-2 demonstrates that with sufficient depth, the model transcends a dependence on supervised learning, developing learned representations from sequence alone that perform equivalently on downstream tasks.

### String-based architectures resolve geometric bias and enable functional prediction for diverse chemistries

Our reliance on SMILES strings is a deliberate design choice to address the limitations of geometric deep learning in this domain. Unlike globular proteins, which fold into stable tertiary structures, therapeutic peptides are often intrinsically disordered or exist as dynamic ensembles. Current 3D methods force these labile molecules into a single static conformation, introducing a geometric bias that misrepresents their behavior in solution. By operating on SMILES, PeptideCLM-2 captures precise chemical connectivity while implicitly allowing the model to learn representations that account for conformational flexibility. This is supported by our results in membrane permeability, where the model successfully inferred 3D-dependent constraints directly from the 1D syntax without requiring explicit structural input.

Notably, PeptideCLM-2 generalizes beyond simple physicochemical descriptors to predict complex biological phenotypes. This capability stems from the fundamental advantage of SMILES-based architectures: the ability to represent a residue not as a fixed label, but as a composite of its constituent chemical parts. While protein language models are structurally limited to the 20 canonical amino acids, our k-mer tokenization strategy allows PeptideCLM-2 to efficiently process the chemical diversity that defines modern peptide therapeutics, natively encoding cyclizations, stereochemical inversions, and diverse chemical modifications. Additionally, this tokenization method allows the model to learn from small molecule chemistry during pretraining and apply that knowledge on downstream tasks.

This structural flexibility not only enabled simple encoding, but also resulted in improved performance over custom architectures and molecular fingerprints on various targets. The model also has demonstrated chemical generalization when successfully predicting the antimicrobial activity of peptides containing noncanonical residues that were held out during training. The finding that embeddings pretrained on chemical syntax (SMILES reconstruction) transfer effectively to functional tasks implies that the chemical features necessary to generalize across molecules are present in the grammar of chemical language.

### Limitations and future directions

While PeptideCLM-2 establishes a new standard for discriminative modeling, several frontiers remain. Peptide aggregation is inherently dynamic, and though our static embeddings predict propensity effectively, capturing true fibrillation kinetics may require training on molecular dynamics trajectories to explicitly model conformational flexibility. Additionally, our reliance on SMILES strings prioritizes topological connectivity over explicit 3D geometry. While the model successfully infers 3D constraints—as evidenced by its permeability predictions—integrating geometric deep learning could further refine performance on highly structure-dependent targets.

Our learning from this work prompts further investigation. While our maximum *k*-mer length of 6 characters was rationally informed, computational constraints limited our evaluation to an atomistic-versus-*k*-mer comparison on our smaller architecture. Future work might explore systematic tokenizer ablations across alternative textual frameworks, including Byte-Pair Encoding (BPE), Subword Peptide Embeddings (SPE), and SELFIES. Furthermore, while our pretraining corpus (ESMAtlas and PubChem) was chosen to represent a wide variety of chemical motifs, a deeper investigation is needed to quantify how representation quality scales as structural targets drift away from the core pretraining data modality. Finally, integrating our unified chemical syntax approach with emerging 3D structural predictors for noncanonical targets, such as AlphaFold 3, represents a promising avenue to mitigate the issues from simple one-dimensional string encoding of high-dimensional and dynamic molecules.

As high-throughput screening^69^ and advanced synthesis^70^ continue to generate large-scale datasets, extending this architecture to the billion-parameter scale promises even greater predictive fidelity. Coupling these predictive oracles with generative frameworks like diffusion models^71^ brings us closer to *de novo* design of noncanonical peptides with precise, multi-parametric profiles.

## Conclusion

We present PeptideCLM-2 as a scalable, open-source chemical language framework designed to bridge the gap between small-molecule cheminformatics and macroscale protein modeling. Rather than relying on rigid, hand-engineered features or localized spatial descriptors, our architecture demonstrates that a unified textual syntax can internalize complex physicochemical dependencies natively from raw SMILES strings. Throughout a rigorous benchmark suite spanning membrane permeability, enzymatic blood stability, and macrocyclic aggregation dynamics, PeptideCLM-2 consistently matched or outperformed specialized architectures and heavily engineered baselines. Its superior performance on out-of-distribution noncanonical challenges confirms that expanding model capacity enables the network to capture transferable, non-linear biophysical principles that generalize beyond the boundaries of natural sequence databases. To foster reproducibility and promote downstream innovation in therapeutic peptide engineering, we release all model weights, tokenizers, and dataset evaluation pipelines to the scientific community.

## Methods

### Dataset curation and preprocessing

To construct a chemically diverse pretraining corpus bridging the gap between small molecules and proteins, we curated data from three primary sources: PubChem,^56^ ESMAtlas,^20^ and the LIPID MAPS Structure Database (LMSD). ^55^

#### Small molecules

We retrieved the complete compound set from PubChem and applied a rigorous filtering cascade to remove non-drug-like entities. Entries were excluded if they were shorter than 20 characters or contained silicon chains. We further removed salts (leading/trailing Br and Cl) and disconnected components (e.g., solvent molecules indicated by ‘.’ within brackets). Polymeric artifacts, specifically repeating silicon oxide motifs (e.g., [Si](=O)[Si](=O)), were identified and discarded. After splitting remaining disconnected components into independent entries and deduplicating the dataset, the final small-molecule corpus contained 108,583,157 unique SMILES strings.

#### Peptides

Peptide sequences were sourced from ESMAtlas. To ensure high-quality structural priors, we filtered for sequences with high predicted confidence (pTM *>* 0.7, pLDDT *>* 0.7) and lengths ≤ 100 amino acids. To reduce redundancy, sequences were clustered at 30% identity using MMseqs2 (--min-seq-id 0.3 -c 0.8 --cov-mode 1), retaining the centroid with the highest product of pTM and pLDDT as the cluster representative. Amino acid sequences were converted to SMILES strings using p2smi.^72^

#### Lipids

All available lipid structures were obtained directly from LMSD.

#### Data Balancing

To prevent model collapse into the dominant small-molecule modality, we employed a balanced sampling strategy during training. Each epoch consisted of an upsampled lipid set (250K), the full peptide set (∼10M), and a random downsampled subset of small molecules (10M), ensuring the model encountered a heterogeneous distribution of chemical syntax.

#### SMILES Processing and Data Augmentation

During self-supervised pretraining, SMILES strings were canonicalized using a random starting atom strategy using ‘doRandom = True‘ in the RDKit MolToSmiles function. This leverages SMILES multiplicity—where a single molecular graph maps uniquely to multiple valid text permutations—as an online data augmentation technique to prevent differences in embeddings from different canonicalizations, as has been seen previously.^73^ For finetuning, SMILES strings were standardized using deterministic, default RDKit canonicalization to ensure reproducible evaluation pipelines. Entries failing RDKit conversion were discarded.

### K-mer SMILES tokenization

Standard atom-level tokenization often results in excessively long sequences for peptide biopolymers, increasing the computational cost of self-attention (O(*n*^2^)). To mitigate this, we developed a custom *k*-mer tokenizer inspired by SMILES Pair Encoding^74^ that compresses peptide text inputs while preserving local chemical semantics. This tokenizer is not equivalent to graph fingerprints such as Morgan or path-based RDKit fingerprints. Those fingerprints enumerate overlapping neighborhoods on the molecular graph, whereas our tokenizer segments a single SMILES string into a linear sequence of substrings consumed sequentially by the transformer encoder.

We first constructed a pre-tokenizer that segments SMILES into atom-level primitives, preserving multi-character elements (e.g., ”Br”, ”Cl”) and stereochemical/charge brackets (e.g., [C@@H], [N+]). We then mined the most frequent contiguous *k*-mers (up to a maximum length of 6 characters) from a reference corpus of 200,000 PubChem small molecules and 200,000 SmProt peptides.^75^ Constraining the vocabulary to a maximum length of 6 ensures that sub-structural fragments (such as localized carbon chains) are encountered with high enough frequency to build robust contextual representations. This candidate list was filtered based on chemical validity and frequency (Table S2), yielding a compact vocabulary of 405 tokens (160 single-atom, 245 *k*-mer). Example vocabulary entries are provided (Table S3). This vocabulary reduces sequence length by approximately 60% compared to character-level encoding, expanding the feasible sequence length boundaries for longer peptide chains.

### Model architecture and pretraining

#### Architecture

PeptideCLM-2 models are trained as BERT-style transformer encoders.^47^ We instantiated three model scales to evaluate performance shifts relative to parameter capacity:

**Small (32M):** 6 layers, 384 hidden dimension, 6 heads.

**Base (114M):** 12 layers, 768 hidden dimension, 12 heads.

**Large (337M):** 24 layers, 1024 hidden dimension, 16 heads.

All models utilize Rotary Positional Embeddings (RoPE) to better capture relative distances in chemical space, along with SwiGLU activation functions and pre-layer normalization for training stability. Full released pretraining hyperparameters can be found (Table S4).

#### Training objectives and Masking Distributions

We employed a hybrid objective function combining self-supervision with property regression:

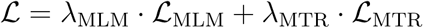

where the objective weights were frozen at *λ*_MLM_ = 0.6 and *λ*_MTR_ = 0.4.

##### Masked language modeling (MLM)

We applied a 25% masking rate using a span-masking strategy to encourage the reconstruction of chemical fragments rather than trivial individual tokens. Span lengths were determined using a Gaussian distribution (*µ* = 3.5, *σ* = 1.0) with a minimum length of one token. For each span, a random starting position was selected, avoiding boundary overruns or overlaps with previously masked regions, and replaced with the [MASK] token ID until the target masking budget was satisfied.

##### Multi-task regression (MTR)

In parallel, a regression head predicted 99 continuous physicochemical descriptors (computed via RDKit, denoted as RDKit99) from the mean-pooled sequence embedding. This MTR head consists of two fully connected layers with SiLU activation. ^52^ All target descriptors were normalized to zero mean and unit variance prior to training. This objective encourages the model to internalize structural boundaries and property vectors during pretraining without requiring explicit descriptor calculations at downstream inference time.

#### Optimization

Models were trained using the AdamW optimizer (*β*_1_ = 0.9*, β*_2_ = 0.98, weight decay 0.01). Pretraining was performed on 8× NVIDIA H100 GPUs with a global batch size of 512 for 3 pseudo-epochs. Each pseudo-epoch consisted of a sample of ∼20.25M molecules, comprising 10M small molecules, the full ESMAtlas peptide corpus, and a 5× upsampling of the LMSD. The learning rate followed a linear warmup for the first 5,000 steps to a peak of 3 × 10*^−^*^4^, followed by cosine annealing to 10% of the peak rate.

### Downstream evaluation protocols

#### Finetuning strategy

For all downstream classification and regression tasks, we replaced the pretraining heads with a task-specific feed-forward network consisting of two hidden layers with GeLU activation and dropout (*p* = 0.1). Optimization was performed using AdamW with an initial learning rate of 1 × 10*^−^*^5^ and a batch size of 16 (Tables S5 and S6).

Train/test splitting protocols were strictly replicated from the benchmark literature for each dataset to preserve fair comparison. The only dataset where prior splits were not available was THPep, where we used random 5-fold cross validation. To rigorously evaluate initialization variance, all downstream fine-tuning and baseline models were trained in triplicate (*n* = 3 independent random seeds), with final performance scores reported as mean ± standard deviation. Models were trained until validation loss plateaued using an early stopping patience and a maximum of 10 epochs.

#### Downstream Baselines and Fingerprint Generation

To evaluate the representative quality of our learned chemical language spaces, we implemented a diverse suite of strong baseline models under identical data splits and evaluation protocols:

- **MorganFP Baseline:** 2048-bit Morgan circular fingerprints were generated using RDKit with a radius of 2 (*r* = 2) to capture topological connectivity fragment counts.
- **RDKit99 Baseline:** 99 continuous continuous physicochemical descriptors mapping property spaces (e.g., LogP, polar surface area, hydrogen bond donor/acceptor counts) were calculated via RDKit. Full descriptor list available (Table S1).
- **Deep Learning Baselines:** We benchmarked against mainstream chemical language frameworks by retraining ChemBERTa-2 (77M MTR) ^37^ and implemented ChemProp^41^ to represent state-of-the-art molecular message-passing graph neural networks.

Tabular fingerprints and descriptors were modeled using an optimized ensemble combining ElasticNet^60^ and ExtraTreesRegressor^61^ in a 50/50 weighting. For specific tasks lacking public splits (e.g., THPep), benchmarks were evaluated across identical triplicate random splits to ensure structural parity.

#### Layer-wise transfer learning, CycPeptMPDB

To assess the intrinsic semantic quality of our raw representations without full weight updates, we frozen the pretrained encoder weights and extracted contextual embeddings from each layer. These layer-wise embeddings were mean-pooled across the sequence dimensions and passed directly to the ElasticNet and ExtraTreesRegressor baseline ensemble to evaluate frozen feature utility. Performance was reported as a function of encoder depth to chart feature optimization across structural transitions.

### Pretraining data overlap with downstream finetuning datasets

To ensure the integrity of our downstream evaluation and quantify potential data leakage from the large-scale pretraining corpus, we performed a structural overlap analysis across all finetuning data splits (Table 2). Downstream sequences were converted to canonicalized SMILES format, which we used to identify the intersection between the target compounds and the pretraining dataset.

**Table 2:** Analysis of unique canonical SMILES structure overlaps with pretraining datasets for finetuning data splits.

| Data Split | Unique SMILES | Overlapping SMILES | Overlap Fraction (%) |
| --- | --- | --- | --- |
| CellPPD Test | 291 | 1 | 0.34% |
| CellPPD Train | 1,065 | 11 | 1.03% |
| THPep | 609 | 4 | 0.66% |
| AmpHGT Test | 5,118 | 132 | 2.58% |
| AmpHGT Train | 9,250 | 268 | 2.90% |
| AmpHGT Val | 2,484 | 78 | 3.14% |
| CycPeptMPDB | 6,649 | 536 | 8.06% |
| PepMSND | 605 | 172 | 28.43% |

Structural redundancy remains low (*<* 3.2%) across the majority of the benchmark splits, including the classification tasks for CellPPD, THPep, and AmpHGT, indicating that down-stream performance on these sets reflects true generalization rather than data memorization. However, a significant structural overlap is observed within the CycPeptMPDB (8.06%) and PepMSND (28.43%) datasets. The elevated redundancy in PepMSND is particularly notable emphasizing the necessity of robust cross-validation partitions when bench-marking self-supervised architectures on these specific endpoints.

## Supporting Information Available

Further data and training parameters are provided in the Supporting Information including: hyperparameter configurations for Small, Base, and Large model variants (Table S1), *k*-mer syntax filtering rules (Table S2), sample token vocabulary examples (Table S3), downstream training and optimization workflows (Table S4), CycPeptMPDB permeability regression statistics (Table S5), classification scores for THPep/CellPPD/AmpHGP/PepMSND (Table S6–S9), PeptideCLM-2 latent space descriptor projections (Figure S1), and layer-wise linear probing permeability profiles (Figure S2).

## Data and Software Availability

All code for pretraining, finetuning, and data processing for this project has been made available at https://github.com/AaronFeller/PeptideCLM-2. Model code and weights for all models are released on HuggingFace at https://huggingface.co/collections/aaronfeller/peptideclm-2. Pretraining data has been released on Zenodo (doi:10.5281/zenodo.17993164) and on Huggingface datasets in the PeptideCLM-2 collection for ease of use. For advancement of modeling therapeutic peptides with noncanonical chemistry, the datasets used for downstream tasks including for membrane permeability, tumor homing, cell penetration, antimicrobial activity, and blood stability have been organized and can be found on the project github.

## Author Contributions

A.L.F. conceived the project, developed the methodology and software, performed the formal analysis and investigation, curated the data, visualized the results, and wrote the original draft. M.S. contributed to the conceptualization and discussions regarding dataset inclusion. S.S. contributed to resources and data curation. C.O.W. and K.D. provided supervision, funding acquisition, and computational resources. All authors reviewed and edited the manuscript.

## Acknowledgement

We would like to express our thanks to Rory Donovan-Maiye and Deepa Korani for their work at Novo Nordisk on large BERT-based modeling of SMILES strings that preceded this project. Novo Nordisk provided compute resources for model pretraining. Finetuning on open classification datasets was performed in part using compute resources provided by NVIDIA. This work was supported in part by NIH grants 1R01 AI148419, 1R01 GM161028, and 1R56 AI179799. C.O.W. was supported by the Blumberg Centennial Professorship in Molecular Evolution.

## TOC Graphic

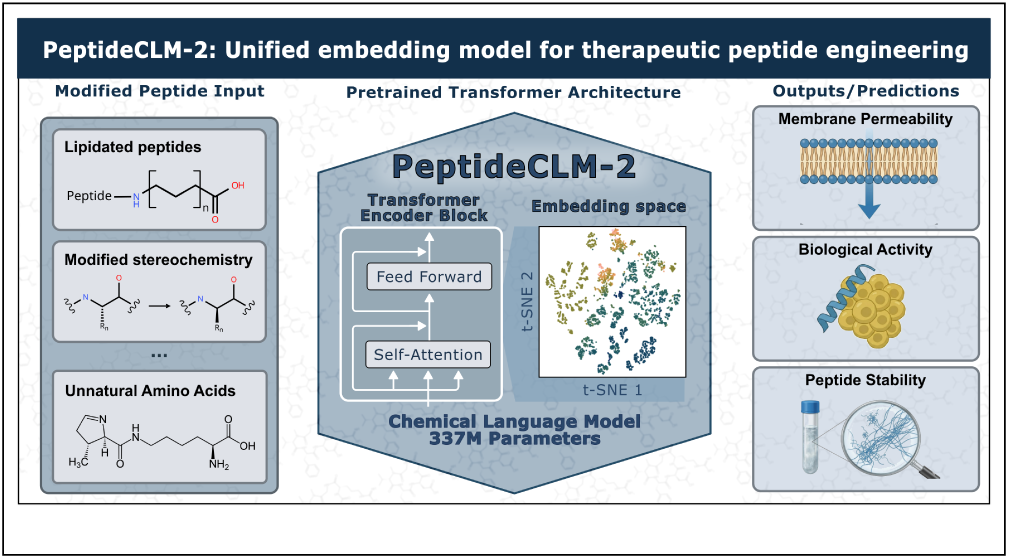

## References

(1) Muttenthaler, M.; King, G. F.; Adams, D. J.; Alewood, P. F. Trends in peptide drug discovery. Nat. Rev. Drug Disc. 2021, 20, 309–325.

(2) Cooper, B. M.; Iegre, J.; O’Donovan, D. H.; Halvarsson, M. O.; Spring, D. R. Peptides as a platform for targeted therapeutics for cancer: peptide–drug conjugates (PDCs). Chem. Soc. Rev. 2021, 50, 1480–1494.

(3) Wang, L.; Wang, N.; Zhang, W.; Cheng, X.; Yan, Z.; Shao, G.; Wang, X.; Wang, R.; Fu, C. Therapeutic peptides: current applications and future directions. Signal Transduct. 2022, 7, 48.

(4) Fetse, J.; Kandel, S.; Mamani, U.-F.; Cheng, K. Recent advances in the development of therapeutic peptides. Trends Pharmacol. Sci. 2023, 44, 425–441.

(5) Barman, P.; Joshi, S.; Sharma, S.; Preet, S.; Sharma, S.; Saini, A. Strategic approaches to improvise peptide drugs as next generation therapeutics. Int. J. Pept. Res. Ther. 2023, 29, 61.

(6) Sharma, K.; Sharma, K. K.; Sharma, A.; Jain, R. Peptide-based drug discovery: Current status and recent advances. Drug Discov. Today 2023, 28, 103464.

(7) Hickey, J. L.; Sindhikara, D.; Zultanski, S. L.; Schultz, D. M. Beyond 20 in the 21st century: prospects and challenges of non-canonical amino acids in peptide drug discovery. ACS Med. Chem. Lett. 2023, 14, 557–565.

(8) Zhang, H.; Chen, S. Cyclic peptide drugs approved in the last two decades (2001–2021). *RSC Chem*. Biol. 2022, 3, 18–31.

(9) Ji, X.; Nielsen, A. L.; Heinis, C. Cyclic Peptides for Drug Development. Angew. Chem. Int. Ed. 2023, 202308251, e202308251.

(10) Lamers, C. Overcoming the shortcomings of peptide-based therapeutics. Future Drug Discov. 2022, 4, FDD75.

(11) Openy, J.; Vega-Ces, S.; Amrahova, G.; Mestdach, E.; Chi, C.; Kissel, B.; ‘t Hart, P. Backbone Alterations in Cyclic Peptides Influence Both Membrane Permeability and Biological Activity. J. Med. Chem. 2025, 68, 24108–24126.

(12) Castro, T. G.; Melle-Franco, M.; Sousa, C. E.; Cavaco-Paulo, A.; Marcos, J. C. Non-canonical amino acids as building blocks for peptidomimetics: Structure, function, and applications. Biomolecules 2023, 13, 981.

(13) Terasaka, N.; Iwane, Y.; Geiermann, A.-S.; Goto, Y.; Suga, H. Recent developments of engineered translational machineries for the incorporation of non-canonical amino acids into polypeptides. International Journal of Molecular Sciences 2015, 16, 6513–6531.

(14) Guo, Z.; Diao, T. Late-Stage Serine Modification Enables Noncanonical Peptide Synthesis. Journal of the American Chemical Society 2025, 147, 33127–33135.

(15) Du, Z.; Caragea, D.; Guo, X.; Li, Y. PepBERT: Lightweight language models for bioactive peptide representation. Future Foods 2026, 13, 100999.

(16) Wang, L.; Pulugurta, R.; Vure, P.; Zhang, Y.; Pal, A.; Chatterjee, P. PepDoRA: A unified peptide language model via weight-decomposed low-rank adaptation. arXiv 2024, 2410.20667.

(17) FernándezDíaz, R.; Ochoa, R.; Hoang, T. L.; Lopez, V.; Shields, D. How to build machine learning models able to extrapolate from standard to modified peptides. J. Cheminform. 2025,

(18) Consortium, T. U. UniProt: the universal protein knowledgebase in 2025. Nucleic Acids Res. 2025, 53, D609–D617.

(19) Rives, A.; Meier, J.; Sercu, T.; Goyal, S.; Lin, Z.; Liu, J.; Guo, D.; Ott, M.; Zitnick, C. L.; Ma, J.; others Biological structure and function emerge from scaling unsupervised learning to 250 million protein sequences. Proc. Natl. Acad. Sci. 2021, 118, e2016239118.

(20) Lin, Z.; Akin, H.; Rao, R.; Hie, B.; Zhu, Z.; Lu, W.; Smetanin, N.; Verkuil, R.; Kabeli, O.; Shmueli, Y.; others Evolutionary-scale prediction of atomic-level protein structure with a language model. Science 2023, 379, 1123–1130.

(21) Hayes, T.; Rao, R.; Akin, H.; Sofroniew, N. J.; Oktay, D.; Lin, Z.; Verkuil, R.; Tran, V. Q.; Deaton, J.; Wiggert, M.; others Simulating 500 million years of evolution with a language model. Science 2025, 387, 850–858.

(22) Elnaggar, A.; Heinzinger, M.; Dallago, C.; Rehawi, G.; Wang, Y.; Jones, L.; Gibbs, T.; Feher, T.; Angerer, C.; Steinegger, M.; others ProtTrans: towards cracking the language of life’s code through self-supervised learning. IEEE Trans. Pattern Anal. Mach. Intell. 2021, 44, 7112–7127.

(23) Alanazi, W.; Meng, D.; Pollastri, G. Porter 6: protein secondary structure prediction by leveraging pre-trained language models (PLMs). Int. J. Mol. Sci. 2024, 26, 130.

(24) Zhang, Z.; Wayment-Steele, H. K.; Brixi, G.; Wang, H.; Kern, D.; Ovchinnikov, S. Protein language models learn evolutionary statistics of interacting sequence motifs. Proc. Natl. Acad. Sci. 2024, 121, e2406285121.

(25) Heinzinger, M.; Littmann, M.; Sillitoe, I.; Bordin, N.; Orengo, C.; Rost, B. Contrastive learning on protein embeddings enlightens midnight zone. NAR Genom. Bioinform. 2022, 4, lqac043.

(26) Sun, Y.; Shen, Y. Structure-informed protein language models are robust predictors for variant effects. Hum. Genet. 2025, 144, 209–225.

(27) Gurev, S.; Youssef, N.; Jain, N.; Marks, D. Variant effect prediction with reliability estimation across priority viruses. bioRxiv 2025, 10.1101/2025.08.04.668549.

(28) Liu, D.; Young, F.; Lamb, K. D.; Claudio Quiros, A.; Pancheva, A.; Miller, C. J.; Macdonald, C.; Robertson, D. L.; Yuan, K. PLM-interact: extending protein language models to predict protein-protein interactions. Nat. Commun. 2025, 16, 9012.

(29) Lu, A. X.; Zhang, H.; Ghassemi, M.; Moses, A. Self-supervised contrastive learning of protein representations by mutual information maximization. BioRxiv 2020, 10.1101/2020.09.04.283929.

(30) Weininger, D. SMILES, a chemical language and information system. J. Chem. Inf. Comput. Sci. 1988, 28, 31–36.

(31) Krenn, M.; Ai, Q.; Barthel, S.; Carson, N.; Frei, A.; Frey, N. C.; Friederich, P.; Gaudin, T.; Gayle, A. A.; Jablonka, K. M.; others SELFIES and the future of molecular string representations. Patterns 2022, 3, 100588.

(32) Wang, S.; Guo, Y.; Wang, Y.; Sun, H.; Huang, J. SMILES-BERT: Large-Scale Unsupervised Pre-training for Molecular Property Prediction. Proc. ACM Conf. Bioinform. Comput. Biol. 2019, 429–436.

(33) Chithrananda, S.; Grand, G.; Ramsundar, B. ChemBERTa: large-scale self-supervised pretraining for molecular property prediction. arXiv 2020, 2010.09885.

(34) Ross, J.; Belgodere, B.; Chenthamarakshan, V.; Padhi, I.; Mroueh, Y.; Das, P. Large-scale chemical language representations capture molecular structure and properties. *Nat*. Mach. Intell. 2022, 4, 1256–1264.

(35) Park, J.-H.; Park, H.; Kim, Y.; Lim, W.; Lee, S. Moleco: Molecular Contrastive Learning with Chemical Language Models for Molecular Property Prediction. Proceedings of the 2024 Conference on Empirical Methods in Natural Language Processing: Industry Track 2024, 408–420.

(36) Park, J.-H.; Kim, Y.; Lee, M.; Park, H.; Lee, S. Moltres: Improving chemical language representation learning for molecular property prediction. arXiv preprint arXiv:2408.01426 2024,

(37) Ahmad, W.; Simon, E.; Chithrananda, S.; Grand, G.; Ramsundar, B. ChemBERTa-2: Towards chemical foundation models. arXiv 2022, 2209.01712.

(38) Lv, L.; Lin, Z.; Li, H.; Liu, Y.; Cui, J.; Chen, C. Y.-C.; Yuan, L.; Tian, Y. ProLLaMA: A protein large language model for multi-task protein language processing. *IEEE Trans*. Artif. Intell. 2025

(39) Mao, Q.; Shang, T.; Xu, W.; Zhai, S.; Zhang, C.; Guo, J.; Su, A.; Li, C.; Duan, H. NCPepFold: Accurate prediction of noncanonical cyclic peptide structures via cyclization optimization with multigranular representation. Journal of Chemical Theory and Computation 2025, 21, 4979–4991.

(40) Zhang, C.; Wang, W.; Zhu, N.; Cao, Z.; Wu, Y.; Mao, Q.; Zhu, C.; Zhang, C.; Guo, J.; Duan, H. AlphaFold3 for noncanonical cyclic peptide modeling: hierarchical bench-marking reveals accuracy and practical guidelines. Journal of Chemical Information and Modeling 2025, 65, 9777–9789.

(41) Heid, E.; Greenman, K. P.; Chung, Y.; Li, S.-C.; Graff, D. E.; Vermeire, F. H.; Wu, H.; Green, W. H.; McGill, C. J. Chemprop: a machine learning package for chemical property prediction. Journal of chemical information and modeling 2023, 64, 9–17.

(42) Burns, J.; Zalte, A.; Green, W. Descriptor-based foundation models for molecular property prediction. *arXiv preprint arXiv:2506.15792* 2025,

(43) Zhang, R.; Wu, H.; Liu, C.; Yang, Q.; Xiu, Y.; Li, K.; Chen, N.; Wang, Y.; Wang, Y.; Gao, X.; others PepLand: a large-scale pre-trained peptide representation model for a comprehensive landscape of both canonical and non-canonical amino acids. Briefings in bioinformatics 2025, 26, bbaf367.

(44) Otero, D. G.; Akbari, O.; Mandapati, A.; Bilodeau, C. PepFoundry: A Pipeline for Building Machine-Learning Ready Representations of Nonstandard Peptides Containing Cycles, Non-natural Residues, Polymer Units, and More. Journal of Chemical Information and Modeling 2026, 66, 1264.

(45) Praski, M.; Adamczyk, J.; Czech, W. Benchmarking pretrained molecular embedding models for molecular representation learning. arXiv 2025, 2508.06199.

(46) Adamczyk, J.; Ludynia, P.; Czech, W. Molecular fingerprints are strong models for peptide function prediction. Bioinformatics 2026, 42, btag179.

(47) Devlin, J.; Chang, M.-W.; Lee, K.; Toutanova, K. BERT: Pre-training of Deep Bidirectional Transformers for Language Understanding. Proc. NAACL 2019, 4171–4186.

(48) Feller, A. L.; Wilke, C. O. Peptide-aware chemical language model successfully predicts membrane diffusion of cyclic peptides. J. Chem. Inf. Model. 2025, 65, 571–579.

(49) Borchani, H.; Varando, G.; Bielza, C.; Larranaga, P. A survey on multi-output regression. WIREs Data Min. Knowl. Discov. 2015, 5, 216–233.

(50) Landrum, G.; others RDKit: Open-source cheminformatics. 2013; http://www.rdkit.org, Online.

(51) Su, J.; Ahmed, M.; Lu, Y.; Pan, S.; Bo, W.; Liu, Y. Roformer: Enhanced transformer with rotary position embedding. Neurocomputing 2024, 568, 127063.

(52) Elfwing, S.; Uchibe, E.; Doya, K. Sigmoid-weighted linear units for neural network function approximation in reinforcement learning. Neural Netw. 2018, 107, 3–11.

(53) Xiong, R.; Yang, Y.; He, D.; Zheng, K.; Zheng, S.; Xing, C.; Zhang, H.; Lan, Y.; Wang, L.; Liu, T. On layer normalization in the transformer architecture. Proc. Int. Conf. Mach. Learn. (ICML) 2020, 10524–10533.

(54) Joshi, M.; Chen, D.; Liu, Y.; Weld, D. S.; Zettlemoyer, L.; Levy, O. SpanBERT: Improving Pre-training by Representing and Predicting Spans. Trans. Assoc. Comput. Linguist. 2020, 8, 64–77.

(55) Sud, M.; Fahy, E.; Cotter, D.; Brown, A.; Dennis, E. A.; Glass, C. K.; Merrill Jr, A. H.; Murphy, R. C.; Raetz, C. R.; Russell, D. W.; others LMSD: LIPID MAPS structure database. Nucleic Acids Res. 2007, 35, D527–D532.

(56) Kim, S.; Chen, J.; Cheng, T.; Gindulyte, A.; He, J.; He, S.; Li, Q.; Shoemaker, B. A.; Thiessen, P. A.; Yu, B.; others PubChem 2025 update. Nucleic Acids Res. 2025, 53, D1516–D1525.

(57) Morgan, H. L. The generation of a unique machine description for chemical structures-a technique developed at Chemical Abstracts Service. J. Chem. Doc. 1965, 5, 107–113.

(58) Ramsundar, B.; Eastman, P.; Walters, P.; Pande, V.; Leswing, K.; Wu, Z. Deep Learning for the Life Sciences: applying deep learning to genomics, microscopy, drug discovery, and more; O’Reilly Media, 2019.

(59) Li, J.; Yanagisawa, K.; Sugita, M.; Fujie, T.; Ohue, M.; Akiyama, Y. CycPeptMPDB: a comprehensive database of membrane permeability of cyclic peptides. J. Chem. Inf. Model. 2023, 63, 2240–2250.

(60) Zou, H.; Hastie, T. Regularization and variable selection via the elastic net. Journal of the Royal Statistical Society Series B: Statistical Methodology 2005, 67, 301–320.

(61) Geurts, P.; Ernst, D.; Wehenkel, L. Extremely randomized trees. Machine learning 2006, 63, 3–42.

(62) Shoombuatong, W.; Schaduangrat, N.; Pratiwi, R.; Nantasenamat, C. THPep: A machine learning-based approach for predicting tumor homing peptides. Comput. Biol. Chem. 2019, 80, 441–451.

(63) Kumar, V.; Agrawal, P.; Kumar, R.; Bhalla, S.; Usmani, S. S.; Varshney, G. C.; Raghava, G. P. Prediction of cell-penetrating potential of modified peptides containing natural and chemically modified residues. Front. Microbiol. 2018, 9, 725.

(64) He, Y.; Song, X.; Wan, H.; Zhao, X. AmpHGT: expanding prediction of antimicrobial activity in peptides containing non-canonical amino acids using multi-view constrained heterogeneous graph transformer. BMC Biol. 2025, 23, 184.

(65) Scodeller, P.; Asciutto, E. K. Targeting tumors using peptides. Molecules 2020, 25, 808.

(66) Yap, C. W. PaDEL-descriptor: An open source software to calculate molecular descriptors and fingerprints. Journal of computational chemistry 2011, 32, 1466–1474.

(67) Hu, H.; Zhang, C.; Xu, Z.; Guo, J.; Su, A.; Li, C.; Duan, H. PepMSND: integrating multi-level feature engineering and comprehensive databases to enhance in vitro/in vivo peptide blood stability prediction. Digital Discovery 2025, 4, 2478–2490.

(68) Liu, Z.; Wang, Y.; Vaidya, S.; Ruehle, F.; Halverson, J.; Soljačić, M.; Hou, T. Y.; Tegmark, M. Kan: Kolmogorov-arnold networks. arXiv preprint arXiv:2404.19756 2024,

(69) Quartararo, A. J.; Gates, Z. P.; Somsen, B. A.; Hartrampf, N.; Ye, X.; Shimada, A.; Kajihara, Y.; Ottmann, C.; Pentelute, B. L. Ultra-large chemical libraries for the discovery of high-affinity peptide binders. Nat. Commun. 2020, 11, 3183.

(70) Niquille, D. L.; Hansen, D. A.; Mori, T.; Fercher, D.; Kries, H.; Hilvert, D. Nonribosomal biosynthesis of backbone-modified peptides. Nat. Chem. 2018, 10, 282–287.

(71) Shin, J.-E.; Riesselman, A. J.; Kollasch, A. W.; McMahon, C.; Simon, E.; Sander, C.; Manglik, A.; Kruse, A. C.; Marks, D. S. Protein design and variant prediction using autoregressive generative models. Nat. Commun. 2021, 12, 2403.

(72) Feller, A. L.; Wilke, C. O. p2smi: A toolkit enabling smiles generation and property analysis for noncanonical and cyclized peptides. Journal of Open Source Software 2025, 10, 8319.

(73) Kosonocky, C. W.; Feller, A. L.; Wilke, C. O.; Ellington, A. D. Using alternative SMILES representations to identify novel functional analogues in chemical similarity vector searches. Patterns 2023, 4.

(74) Li, X.; Fourches, D. SMILES pair encoding: a data-driven substructure tokenization algorithm for deep learning. J. Chem. Inf. Model. 2021, 61, 1560–1569.

(75) Li, Y.; Zhou, H.; Chen, X.; Zheng, Y.; Kang, Q.; Hao, D.; Zhang, L.; Song, T.; Luo, H.; Hao, Y.; others SmProt: a reliable repository with comprehensive annotation of small proteins identified from ribosome profiling. Genomics Proteomics Bioinformatics 2021, 19, 602–610.

