## Supporting Figures and Tables for "Scaling SMILES-based chemical language models for therapeutic peptide engineering"

### Contents

**Supplementary Tables** **S2**

S1: *RDKit Descriptors by Type* . . . . . S2
S2: *Kmer Filtering Rules* . . . . . S3
S3: *Sample Token Vocabulary* . . . . . S3
S4: *Pretraining Parameters* . . . . . S4
S5: *Downstream Workflow Settings* . . . . . S5
S6: *Finetuning Regularization Mappings* . . . . . S6
S7: *Permeability Benchmark Statistics* . . . . . S6
S8: *Tumor-Homing Benchmark Metrics* . . . . . S7
S9: *Cell-Penetration Benchmark Metrics* . . . . . S7
S10: *Antimicrobial Benchmark Metrics* . . . . . S7
S11: *PepMSND Blood Stability Statistics* . . . . . S7

**Supplementary Figures** **S8**

S1: *t-SNE Embeddings Visualizations* . . . . . S8
S2: *Layer Probing Profiles* . . . . . S9

### Supplementary Tables

Table S1: RDKit Descriptors Organized by Type

| Descriptor Type | Descriptors |
|---|---|
| **Topological Indices** | Chi0, Chi0n, Chi0v, Chi1, Chi1n, Chi1v, Chi2n, Chi2v |
| **Morgan Fingerprints** | FpDensityMorgan1, FpDensityMorgan2, FpDensityMorgan3 |
| **Kappa Indices** | Kappa1, Kappa2, Kappa3 |
| **Molecular Properties** | ExactMolWt, MaxAbsPartialCharge, MaxPartialCharge, MinAbsPartialCharge, MinPartialCharge, MolLogP, MolMR, MolWt |
| **Structural Counts** | RingCount, HeavyAtomCount, HeavyAtomMolWt, FractionCSP3 |
| **SA Descriptor** | HallKierAlpha, LabuteASA, TPSA |
| **Atom Count Descriptors** | NHOHCount, NOCount, NumHAcceptors, NumHDonors, NumHeteroatoms |
| **Ring Counts** | NumAliphaticCarbocycles, NumAliphaticHeterocycles, NumAliphaticRings, NumAromaticCarbocycles, NumAromaticHeterocycles, NumAromaticRings |
| **Electronic Descriptors** | NumRadicalElectrons, NumValenceElectrons |
| **Rotatable Bonds** | NumRotatableBonds |
| **Saturated Structure Counts** | NumSaturatedCarbocycles, NumSaturatedHeterocycles, NumSaturatedRings |
| **PEOE Descriptors** | PEOE_VSA1, PEOE_VSA2, PEOE_VSA3, PEOE_VSA4, PEOE_VSA5, PEOE_VSA6, PEOE_VSA7, PEOE_VSA8, PEOE_VSA9, PEOE_VSA10, PEOE_VSA11, PEOE_VSA12, PEOE_VSA13, PEOE_VSA14 |
| **SMR Descriptors** | SMR_VSA1, SMR_VSA2, SMR_VSA3, SMR_VSA4, SMR_VSA5, SMR_VSA6, SMR_VSA7, SMR_VSA8, SMR_VSA9, SMR_VSA10 |
| **SlogP Descriptors** | SlogP_VSA1, SlogP_VSA2, SlogP_VSA3, SlogP_VSA4, SlogP_VSA5, SlogP_VSA6, SlogP_VSA7, SlogP_VSA8, SlogP_VSA9, SlogP_VSA10, SlogP_VSA11, SlogP_VSA12 |
| **Functional Groups** | fr_amide, fr_NH0, fr_NH1, fr_NH2, fr_COO, fr_priamide, fr_guanido, fr_imidazole, fr_phenol, fr_Al_OH, fr_C_O, fr_ether, fr_alkyl_halide, fr_unbrch_alkane, fr_aryl_methyl, fr_benzene, fr_ester, fr_ketone, fr_methoxy, fr_sulfide, fr_sulfonamd |

Table S2: Filtering rules applied during construction of the kmer vocabulary.

| # | **Remove tokens:** |
|---|---|
| 1 | with numbers |
| 2 | that start with `')'` |
| 3 | that end with `'('` |
| 4 | that contain `')*'` w/o a leading `'*('` |
| 5 | with more than 4 atom characters |
| 6 | with fewer than 1,000 occurrences |

Table S3: Sample tokens extracted from the initialized kmer tokenizer.

| **Tokens** |
|---|
| =NC, #Cc, c(CO), n(C)c, n(C), CN=C, (CCO), CCO), C(C)N, C#C, C(CC), CS(=O), NC(C), CSc, NN, CC(O), CNc, CCn |

Table S4: Pretraining model configurations and hardware optimization mappings.

| Setting | Value |
|---|---|
| Architecture | BERT-style encoder (Small/Base/Large) |
| Parameter scales | 32M, 114M, 337M |
| Small architecture | 14 blocks, embed 512, 8 heads, FFN 768 |
| Base architecture | 24 blocks, embed 768, 12 heads, FFN 1024 |
| Large architecture | 32 blocks, embed 1024, 16 heads, FFN 2048 |
| Tokenizer | kmer SMILES (vocab: 405) |
| Max sequence length | 2048 |
| Batch size | 512 seqs (global) |
| Optimizer | AdamW |
| AdamW $(\beta_1, \beta_2, \epsilon)$ | $(0.9, 0.98, 1\text{e}{-8})$ |
| Weight decay | 0.01 |
| Learning rate (peak) | 3e−4 |
| LR schedule | Cosine decay with linear warmup |
| Warmup steps | 5,000 |
| Training steps | 100k |
| Dropout (attn/ffn) | 0.1 / 0.1 |
| Masking rate (MLM) | 25% |
| Span masking | $\mathcal{N}(\mu=3.5, \sigma=1)$ spans |
| MTR heads | 2-layer MLP, SiLU, mean-pooled embedding |
| Regression targets | 99 RDKit descriptors (normalized) |
| Loss weights | $\lambda_{\text{MLM}} = 0.6,\ \lambda_{\text{MTR}} = 0.4$ |
| Precision | bfloat16 |
| Gradient clipping | 0.1 |
| Gradient accumulation | 1 / 2 / 4 (small/base/large) |
| Data aug. | RDKit randomized SMILES |

Table S5: Released downstream training settings organized by validation workflow.

| Workflow | Released setting |
|---|---|
| CycPeptMPDB regression | Mean-pooled transformer embeddings with a small regression head; MSE loss; AdamW; batch size 16; max 10 epochs; validation every 0.2 epoch; early stopping on validation loss; outer held-out folds with inner-fold ensembling. |
| Classification benchmarks | Rank-16 LoRA adapters on the attention projection layers with a SiLU projection head; AdamW; default batch size 16; linear warmup scheduler; max 10 epochs; early stopping patience 5; fixed splits when available and five-fold CV otherwise. |
| ChemBERTa baseline | Batch size 8; learning rate $2 \times 10^{-5}$; max 8 epochs; early stopping patience 2. |
| PepMSND | Mean-pooled transformer embedding fused with one-hot Species/Environment covariates; BCEWithLogitsLoss; AdamW with learning rate $1 \times 10^{-5}$; batch size 32; max 10 epochs; early stopping patience 3. |
| Reported summaries | Classification metrics: MCC, AUROC, F1. Regression metrics: $R^2$, RMSE, MAE. Publication figures aggregate repeated runs across three random seeds unless otherwise noted. |

Table S6: Finetuning optimization criteria and validation dimensions.

| Setting | Value (Small, Base, Large) |
|---|---|
| Task heads | Regression |
| Loss | MSE |
| Batch size | 16 |
| Learning rate | 3e−4 |
| Optimizer | AdamW |
| Weight decay | 0.01 |
| LR schedule | No decay; no warmup |
| Dropout (head) | 0.1 |
| Max epochs (per fold) | 10 |
| Evaluation | 20% epoch (early stopping patience 3) |
| CV scheme | Outer holdout + inner K-fold |
| Ensembling | Mean of checkpoints across inner folds |
| Class imbalance | Equal sampling across bins |
| Input length | No truncations required |
| Replicates | 3 (report mean ± std) |

Table S7: CycPeptMPDB test-set regression metrics from the released triplicate permeability runs, reported as mean ± SD.

| Size / Control | Released setting | $R^2$ | RMSE | MAE |
|---|---|---|---|---|
| Control | Random initialization | $-0.079 \pm 0.048$ | $0.805 \pm 0.018$ | $0.622 \pm 0.010$ |
| **Small** | MLM | $0.237 \pm 0.140$ | $0.675 \pm 0.061$ | $0.516 \pm 0.052$ |
| | Hybrid | $0.232 \pm 0.095$ | $0.678 \pm 0.042$ | $0.527 \pm 0.037$ |
| | **MTR** | $\mathbf{0.378 \pm 0.048}$ | $\mathbf{0.611 \pm 0.024}$ | $\mathbf{0.467 \pm 0.025}$ |
| **Base** | MLM | $0.489 \pm 0.086$ | $0.553 \pm 0.048$ | $0.420 \pm 0.037$ |
| | Hybrid | $0.495 \pm 0.042$ | $0.550 \pm 0.023$ | $0.419 \pm 0.017$ |
| | **MTR** | $\mathbf{0.502 \pm 0.034}$ | $\mathbf{0.547 \pm 0.019}$ | $\mathbf{0.412 \pm 0.017}$ |
| **Large** | MLM | $0.582 \pm 0.041$ | $0.500 \pm 0.025$ | $0.374 \pm 0.018$ |
| | **Hybrid** | $\mathbf{0.613 \pm 0.044}$ | $\mathbf{0.482 \pm 0.028}$ | $\mathbf{0.361 \pm 0.022}$ |
| | MTR | $0.582 \pm 0.018$ | $0.501 \pm 0.011$ | $0.374 \pm 0.010$ |

Table S8: THPep tumor-homing test-set metrics across three random seeds for the released benchmark models, reported as mean ± SD.

| Model | MCC | AUROC | F1 |
|---|---|---|---|
| RDKit99 | $0.673 \pm 0.092$ | $0.932 \pm 0.029$ | $0.766 \pm 0.066$ |
| MorganFP | $0.597 \pm 0.040$ | $0.903 \pm 0.009$ | $0.707 \pm 0.027$ |
| ChemBERTa-2 | $0.618 \pm 0.118$ | $0.921 \pm 0.013$ | $0.719 \pm 0.105$ |
| CheMeleon | $0.600 \pm 0.062$ | $0.888 \pm 0.049$ | $0.703 \pm 0.051$ |
| PeptideCLM-2 (MLM) | $\mathbf{0.756 \pm 0.019}$ | $\mathbf{0.949 \pm 0.006}$ | $\mathbf{0.826 \pm 0.012}$ |
| PeptideCLM-2 (Hybrid) | $0.747 \pm 0.036$ | $0.940 \pm 0.019$ | $0.818 \pm 0.022$ |
| PeptideCLM-2 (MTR) | $0.698 \pm 0.036$ | $0.924 \pm 0.016$ | $0.784 \pm 0.029$ |

Table S9: CellPPD-Mod cell-penetrating test-set metrics across three random seeds for the released benchmark models, reported as mean ± SD.

| Model | MCC | AUROC | F1 |
|---|---|---|---|
| RDKit99 | $0.829 \pm 0.010$ | $0.968 \pm 0.001$ | $0.915 \pm 0.005$ |
| MorganFP | $0.891 \pm 0.003$ | $\mathbf{0.992 \pm 0.000}$ | $0.942 \pm 0.002$ |
| ChemBERTa-2 | $0.768 \pm 0.008$ | $0.943 \pm 0.004$ | $0.876 \pm 0.004$ |
| CheMeleon | $\mathbf{0.892 \pm 0.016}$ | $0.983 \pm 0.000$ | $\mathbf{0.945 \pm 0.007}$ |
| PeptideCLM-2 (MLM) | $0.875 \pm 0.004$ | $0.981 \pm 0.002$ | $0.935 \pm 0.003$ |
| PeptideCLM-2 (Hybrid) | $0.860 \pm 0.003$ | $0.976 \pm 0.001$ | $0.926 \pm 0.003$ |
| PeptideCLM-2 (MTR) | $0.867 \pm 0.004$ | $0.975 \pm 0.001$ | $0.929 \pm 0.002$ |

Table S10: AMP-HGT antimicrobial test-set metrics across three random seeds for the released benchmark models, reported as mean ± SD.

| Model | MCC | AUROC | F1 |
|---|---|---|---|
| RDKit99 | $0.835 \pm 0.007$ | $0.967 \pm 0.002$ | $0.923 \pm 0.003$ |
| MorganFP | $0.836 \pm 0.005$ | $\mathbf{0.986 \pm 0.001}$ | $0.923 \pm 0.002$ |
| ChemBERTa-2 | $0.814 \pm 0.010$ | $0.957 \pm 0.007$ | $0.914 \pm 0.004$ |
| CheMeleon | $0.818 \pm 0.009$ | $0.965 \pm 0.002$ | $0.916 \pm 0.004$ |
| PeptideCLM-2 (MLM) | $\mathbf{0.884 \pm 0.013}$ | $0.980 \pm 0.004$ | $\mathbf{0.946 \pm 0.006}$ |
| PeptideCLM-2 (Hybrid) | $0.850 \pm 0.014$ | $0.972 \pm 0.003$ | $0.930 \pm 0.007$ |
| PeptideCLM-2 (MTR) | $0.853 \pm 0.039$ | $0.972 \pm 0.011$ | $0.931 \pm 0.018$ |

Table S11: PepMSND test-set metrics from the released triplicate prediction exports, reported as mean ± SD across runs.

| Released model | MCC | AUROC | F1 |
|---|---|---|---|
| MLM-MTR embedding + linear head | $0.419 \pm 0.026$ | $0.791 \pm 0.007$ | $0.598 \pm 0.039$ |
| MLM-MTR embedding + KAN head | $\mathbf{0.640 \pm 0.016}$ | $\mathbf{0.879 \pm 0.002}$ | $\mathbf{0.788 \pm 0.008}$ |

### Supplementary Figures

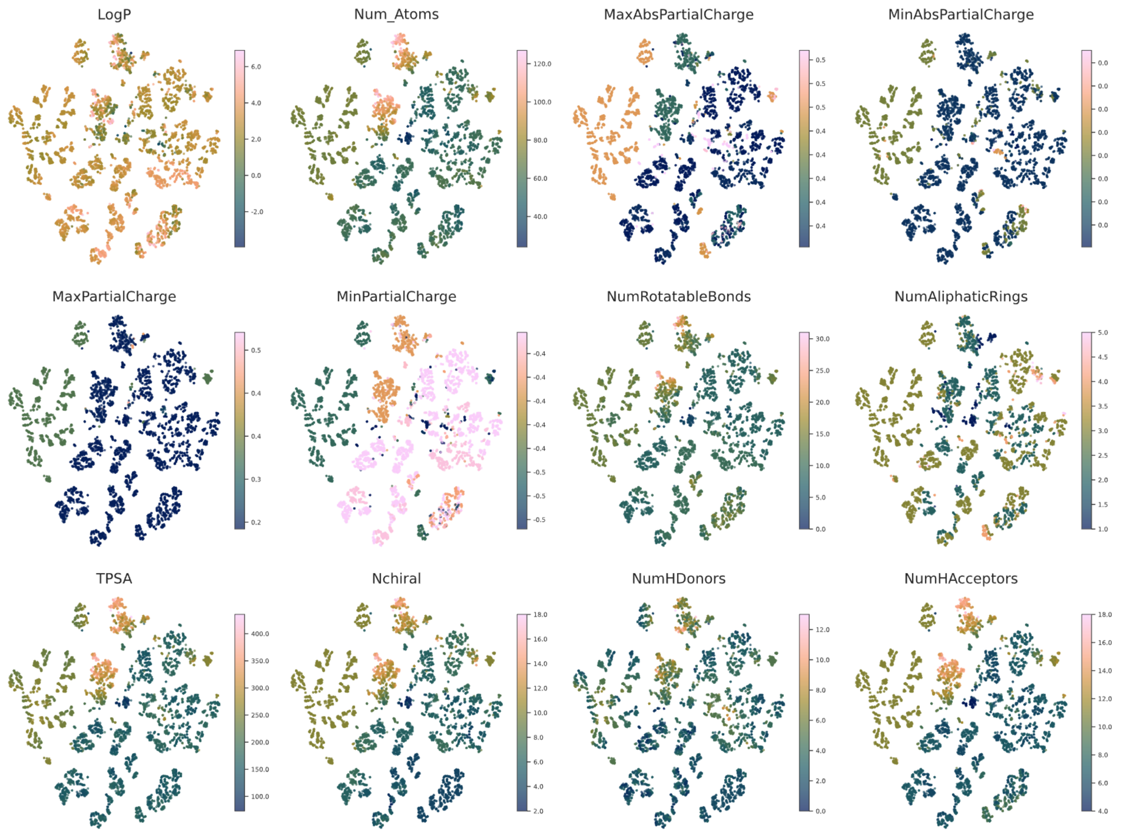


Figure S1: **Latent space visualization of key molecular descriptors.** 2D t-SNE plots of PeptideMLM embeddings (large model) displayed with colorbars of with calculated physicochemical properties. Clustering demonstrates the correlation between the learned embedding space and specific chemical features including physical and chemical properties.

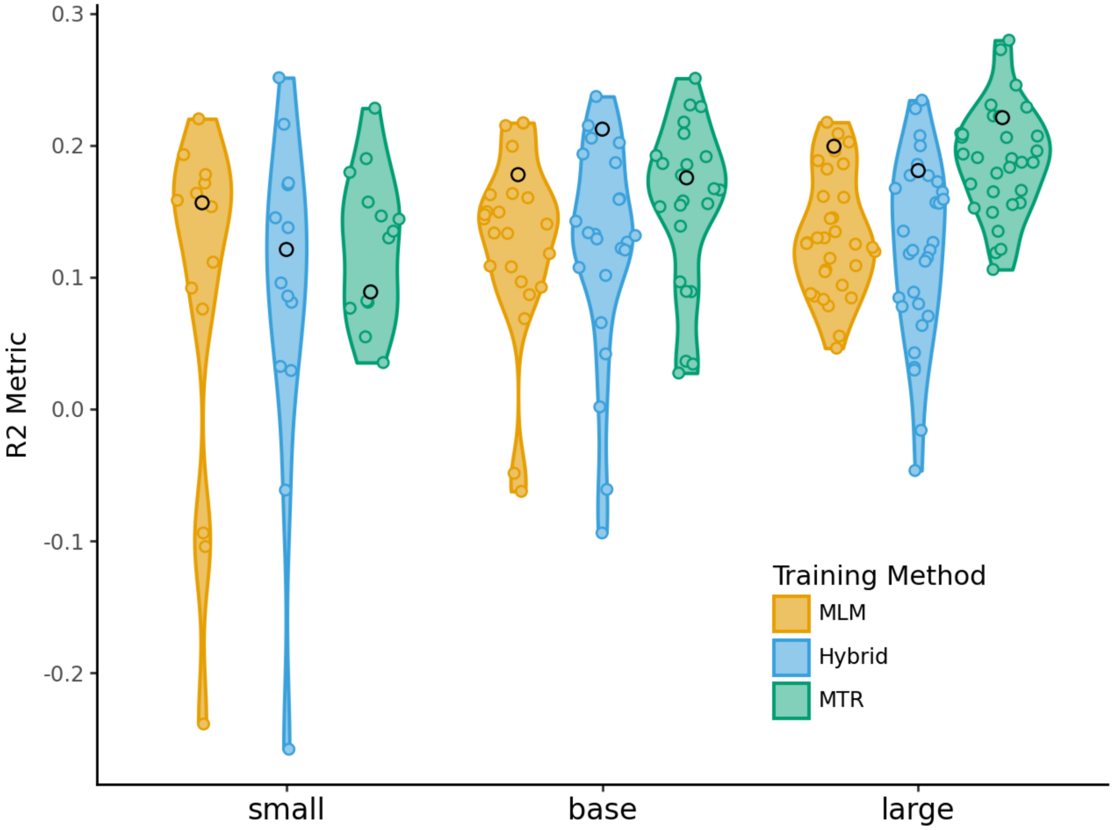


Figure S2: **Linear probing of embeddings for PAMPA permeability.** Violin plots displaying the distribution of $R^2$ scores obtained via LassoCV regression on the CycPeptMPDB dataset. Models were trained using mean-pooled embeddings extracted from pre-trained encoders, stratified by architecture size (Small, Base, Large) and training objective (MLM, Hybrid, MTR). The spread within each violin represents the variance in predictive power across embeddings extracted from different model layers. The final layer is indicated with a black dot.
